# Agromonas: a rapid disease assay for *Pseudomonas syringae* growth in agroinfiltrated leaves

**DOI:** 10.1101/2020.08.10.243808

**Authors:** Pierre Buscaill, Nattapong Sanguankiattichai, Yoon Joo Lee, Jiorgos Kourelis, Gail Preston, Renier A. L. van der Hoorn

## Abstract

The lengthy process to generate transformed plants is a limitation in current research on the interactions of the model plant pathogen *Pseudomonas syringae* with plant hosts. Here we present an easy method called agromonas, where we quantify *P. syringae* growth in agroinfiltrated leaves of *Nicotiana benthamiana* using a cocktail of antibiotics to select *P. syringae* on plates. As a proof of concept, we demonstrate that transient expression of PAMP receptors reduces bacterial growth and that transient depletion of a host immune gene and transient expression of a T3 effector increase *P. syringae* growth in agromonas assays. We show that we can rapidly achieve structure-function analysis of immune components and test the function of immune hydrolases. The agromonas method is easy, fast and robust for routine disease assays with various *Pseudomonas* strains without transforming plants or bacteria. The agromonas assay offers reliable opportunity for further comprehensive analysis of plant immunity.

**One sentence summary:** Agromonas is a rapid and robust disease assay to monitor *Pseudomonas syringae* growth in agroinfiltrated leaves expressing immune components and their suppressors.

## INTRODUCTION

Understanding the plant immune system and microbial pathogenicity is essential to improve plant biotechnologies and crop protection. To evaluate the level of resistance of a plant or the virulence of bacterial pathogens, the routine method relies on infection assays that quantify bacterial growth (*i.e.* colony count assays). Colony count assays are usually performed on stable transformant plants. However, generation of stable transgenic lines is time and resource consuming and is limited to plant species that are amenable to genetic transformation. Therefore, there is a need for faster disease assays particularly in the studies of the model plant pathogen *Pseudomonas syringae*, which causes important economic damages in many plant species.

Rapid overexpression and transcript depletion of various exogenous and endogenous genes is facilitated by *Agrobacterium tumefaciens*-mediated transient expression (agroinfiltration). Agroinfiltration is used throughout plant science to study protein localisation and for their biochemical characterisation. Agroinfiltrated leaves are routinely used to study the interaction between the model plant *Nicotiana benthamiana* and the potato blight pathogen *Phytophthora infestans* (Dagdas *et al.*, 2018; Bozkurt *et al.*, 2014; Chaparro-Garcia *et al.*, 2011) and other *Phytophthora* species. However, agroinfiltrated *N. benthamiana* leaves are not routinely used for disease assays with *P. syringae*. One problem is that selective isolation of *P. syringae* from agroinfiltrated tissue is challenging because there is an overlap of endogenous or introduced antibiotics resistance. For example, rifampicin is commonly used to select for antibiotic resistance genes in the genome of *A. tumefaciens* and *P. syringae* strains, whereas resistance to kanamycin is frequently used to maintain plasmids in both bacteria and therefore these antibiotics are not useful for selective isolation in co-inoculated tissue.

A solution to this problem is to use selection for bacterial endogenous resistance to antibiotics. The combination of 10 µg/ml cetrimide, 10 µg/ml fucidin and 50µg/ml cephaloridine (CFC, Figure 1a) permits the selection of *Pseudomonas* species (Mead and Adams, 1977). CFC is used for the selective isolation of *Pseudomonas* species during the microbiological examination of environmental, clinical, food and plant samples (Krueger and Sheikh 1987; Hill *et al.*, 2005; Pantazi *et al.*, 2008; Fones *et al.*, 2010; Straub *et al.*, 2018), but has not yet been exploited in combination with agroinfiltrated leaves.

**Figure 1.**
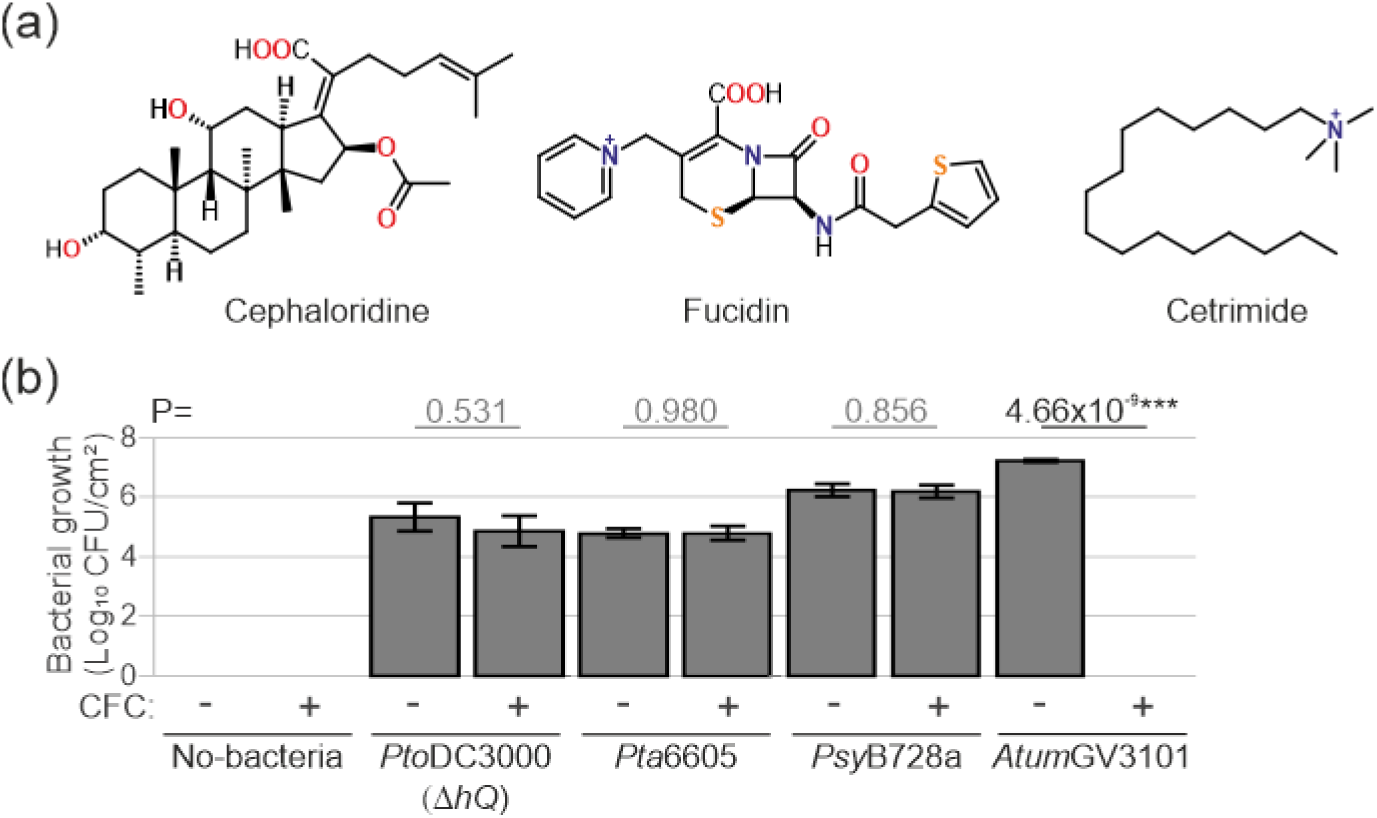
Selection against *Agrobacterium tumefaciens*. (a) Chemical structures of cephaloridine, fucidin and cetrimide (CFC). (b) *P. syringae* grows on CFC selection, *Agrobacterium* does not. *N. benthamiana* leaves were infiltrated with 1×10^6^ CFU/ml *P. syringae* or 1×10^8^ CFU/ml *Atum*GV3101 and bacterial populations were determined three days later using colony count method using LB plates containing CFC or not. Error bars indicate SE of n=3 biological replicates. Student’s *t*-test statistics (*** p<0.001). CFU, colony-forming units.

Here, we use CFC selection to establish a rapid and easy disease assay to quantify growth of *P. syringae* from agroinfiltrated leaves. We tested whether immunity can be studied in agroinfiltrated leaves even though these leaves contain *A. tumefaciens*. The method we developed is called ‘<u>agromonas</u>’ because is based on <u>agro</u>infiltration followed by inoculation of *Pseudo<u>monas</u> syringae* by both infiltration and spray inoculation. We demonstrate that the agromonas assay can be applied to different *P. syringae* strains and demonstrate its adaptability to study the impact of immune components and bacterial effectors on *P. syringae* growth *in planta*.

## RESULTS

### CFC facilitates *P. syringae* selection from agroinfiltrated tissues

To confirm that CFC facilitates *P. syringae* selection we tested three different *P. syringae* strains that are pathogenic on *N. benthamiana*. We tested *P. syringae* pv. *tomato* DC3000, the causative agent of the bacterial speck disease of tomato lacking the type III effector gene *hopQ1-1*[*Pto*DC3000(∆*hQ*)]; *P. syringae* pv. *tabaci* 6605 (*Pta*6605), the causative agent for wildfire disease in tobacco; and *P. syringae* pv. *syringae* B728a (*Psy*B728a), the causative agent of bacterial brown spot of bean. We also included *A. tumefaciens* GV3101 (*Atum*GV3101) the non-oncogenic strain that is routinely used for agroinfiltration. These strains were infiltrated into *N. benthamiana* leaves, and at three days post-infiltration (3 dpi), leaf extracts were generated, diluted in water and plated out on LB medium with or without CFC selection. All tested *P. syringae* pathovars grew equally well on LB medium supplemented with or without CFC (Figure 1b), demonstrating that CFC does not affect *P. syringae* growth. By contrast, *Atum*GV3101 did not grow at all on plates containing CFC (Figure 1b). Likewise, CFC also blocks growth of the non-oncogenic *Atum*C58C1 as well as all tested *Xanthomonas* strains (Table S1), confirming the selectivity of CFC.

To apply CFC selection to facilitate the selective isolation of *P. syringae* from agroinfiltrated leaves, *N. benthamiana* leaves were first infiltrated with *Atum*GV3101. Two days later, each of the three different *P. syringae* strains were infiltrated into the agroinfiltrated regions. Three days later, leaf homogenates were plated onto LB plates containing CFC or gentamicin (Figure 2a). While *P. syringae* strains were specifically isolated on CFC plates, *Atum*GV3101 was isolated on plates containing gentamicin (Figure 2b,c). No colonies with Agrobacterium morphology were detected on CFC plates (Figure 2b), demonstrating that *A. tumefaciens* cannot grow on CFC plates, even when *P. syringae* is growing. These data demonstrate that CFC-containing medium facilitates the selection of living *Pseudomonas* spp. from agroinfiltrated leaves.

**Figure 2.**
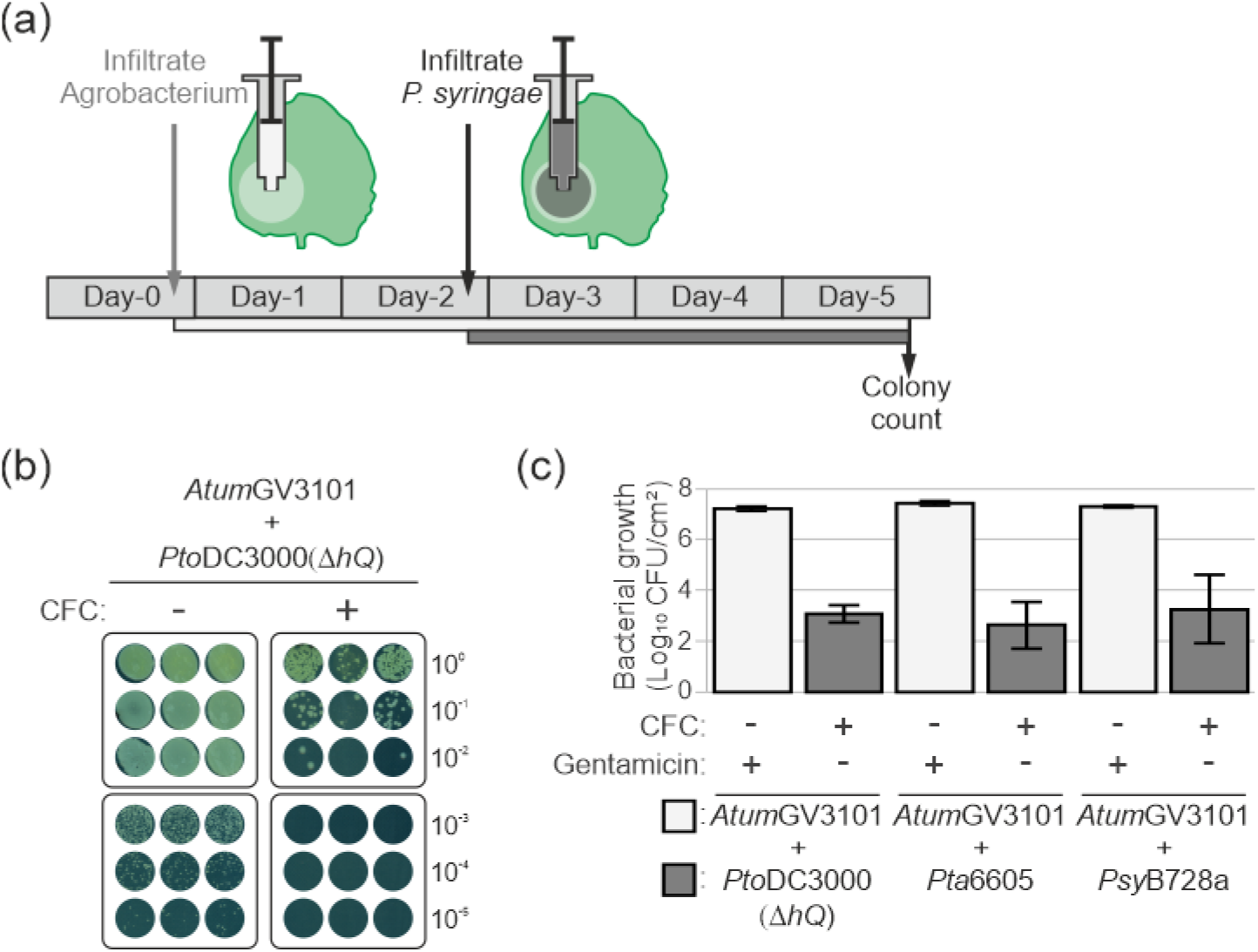
Concept of agromonas assay. (a) Experimental procedure for agromonas assay. Two days after agroinfiltration, agroinfiltrated leaves are infiltrated with *P. syringae* bacteria. Bacterial growth is measured three days later by a classic colony count on LB agar plates containing CFC. (b) CFC selects *P. syringae* from agroinfiltrated leaves. Agroinfiltrated leaves were infiltrated with 1×10^6^ CFU/ml *Pto*DC3000(∆*hQ*) and three days later leaf extracts were diluted, and each dilution were plated onto medium supplemented with or without CFC. Pictures were taken 48 h later. (c) Selective isolation of *Pseudomonas* spp. from agroinfiltrated leaves. Agroinfiltrated leaves were infiltrated with 1×10^6^ CFU/ml *P. syringae* and three days later, leaf extracts were plated on medium containing CFC or gentamicin to select *P. syringae* or *Agrobacterium*, respectively. Error bars indicate SE of n=3 biological replicates.

While comparing unmixed infection with mixed infection (Figures 1b and 2c), we observed that the presence of *A. tumefaciens* suppresses *P. syringae* growth, as previously reported (Rico *et al.*, 2010). We compared the growth of *Pto*DC3000(∆*hQ*) in the presence and absence of *Atum*GV3101*. Pto*DC3000(∆*hQ*) grew 9-fold less in *N. benthamiana* leaves infiltrated with *Atum*GV3101 as compared to a non-agroinfiltrated sample (Figure S1), confirming that *A. tumefaciens* reduces *P. syringae* growth *in planta*. Therefore, it is essential to use agroinfiltrated leaves expressing the empty vector (EV) as control in agromonas assays.

### PAMP receptors reduce bacterial growth in agromonas assay

To demonstrate that the agromonas assay can be used to study genes that confer immunity, we used two PRRs (pattern recognition receptors) that are not present in *N. benthamiana*. We tested tomato FLS3 (flagellin-sensing 3) and Arabidopsis EFR (EF-Tu receptor) which recognize the flgII-28 epitope of flagellin and the elf18 epitope of EF-Tu, respectively (Hind *et al.*, 2016; Cai *et al.*, 2011; Kunze *et al.*, 2004; Zipfel *et al.*, 2006).

To confirm the functionality of FLS3 and EFR upon agroinfiltration, we transiently expressed FLS3 and EFR in *N. benthamiana* leaves and measured the production of reactive oxygen species (ROS) upon treatment with flgII-28 and elf18. Leaves transiently expressing FLS3 were able to release a ROS burst upon flgII-28 treatment, whereas EFR and EV expressing leaves remained unresponsive to flgII-28 (Figure 3a). Likewise, leaves that transiently express EFR were able to release an oxidative burst upon elf18 treatment, whereas FLS3 and EV expressing leaves remained unresponsive to elf18 (Figure 3a). These results show that FLS3 and EFR are functional in *N. benthamiana*, consistent with previous studies (Hind *et al.*, 2016; Lacombe *et al.*, 2010).

**Figure 3.**
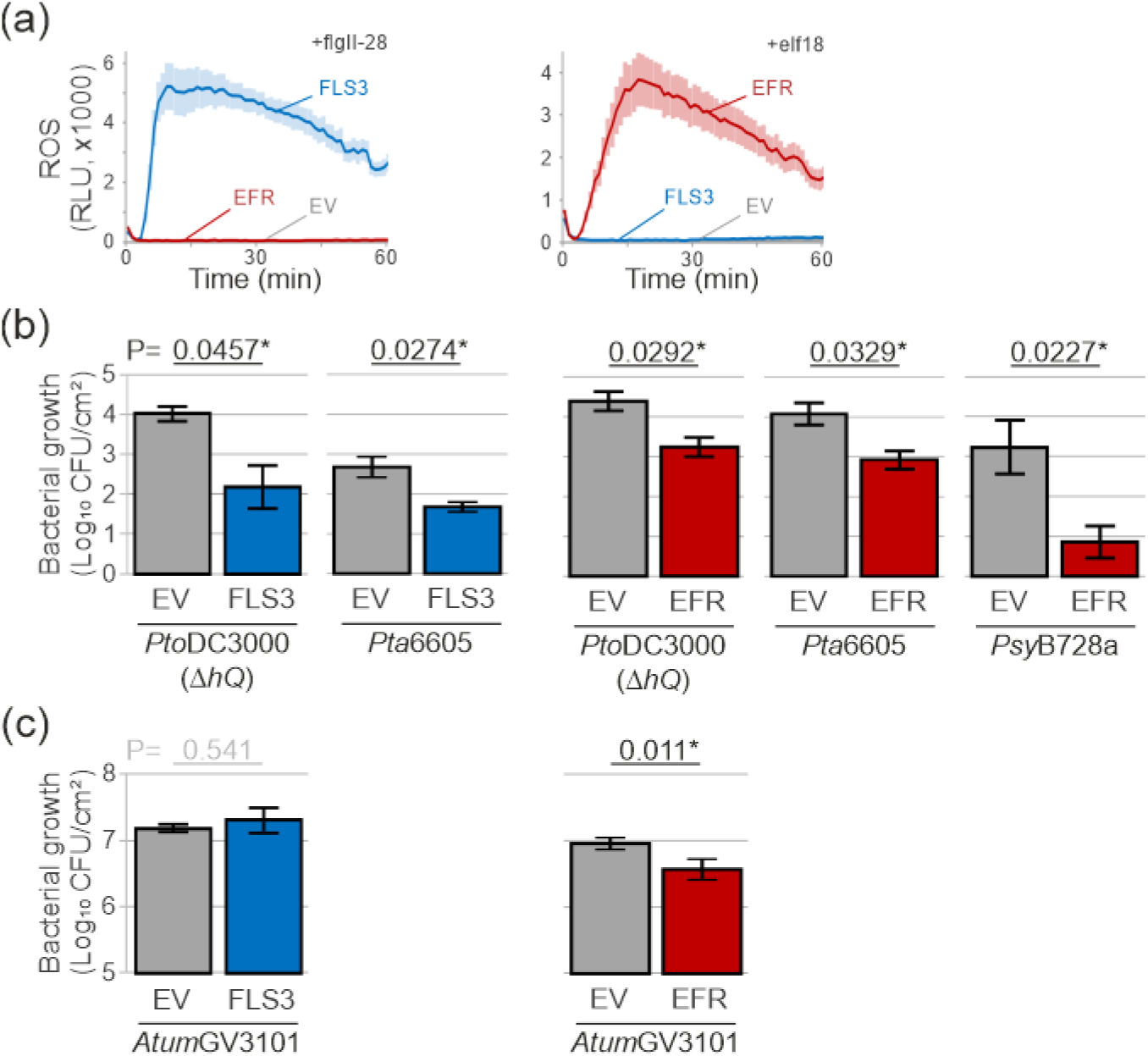
PAMP receptors reduce bacterial growth in the agromonas assay. (a) Transient expression of tomato FLS3 and Arabidopsis EFR in *N. benthamiana* confers flgII-28 and elf18 responsiveness, respectively. Leaf discs from agroinfiltrated leaves expressing FLS3 (blue), EFR (red) or EV (grey) were treated with 100 nM flgII-28 or elf18 and ROS was measured in relative light units (RLU). Error intervals (shaded regions) indicate SE of n=12 biological replicates. (b) Transient expression of FLS3 or EFR reduces *P. syringae* growth. Two days after agroinfiltration, agroinfiltrated leaves expressing FLS3 (blue), EFR (red) or EV (grey) were spray-inoculated with the indicated strains of *P. syringae* (at 1×10^8^ CFU/ml) and bacterial growth was measured three days later using CFC selection. Error bars indicate SE of n=3 biological replicates. Student’s *t*-test statistics (* p<0.05). (c) Transient expression of EFR, but not FLS3, affect Agrobacterium growth. Bacterial growth of *Atum*GV3101 was measured by plating the leaf extracts described in (b) on medium containing gentamicin. Error bars indicate SE of n=3 replicates. Student’s *t*-test statistics (* p<0.05).

We next tested whether agroinfiltrated leaves expressing FLS3 have enhanced resistance to *P. syringae* upon infection. Agroinfiltrated leaves of *N. benthamiana* plants transiently expressing FLS3 showed reduced bacterial growth of both *Pto*DC3000(∆*hQ*) and *Pta*6605 strains compared to leaves expressing the EV control (Figure 3b). Similarly, EFR transient expression caused a strong reduction in the growth of *Pto*DC3000(∆*hQ*), *Pta*6605 and *Psy*B728a (Figure 3b). Altogether these data demonstrate that agroinfiltration of FLS3 and EFR increases immunity to *P. syringae* in *N. benthamiana*.

We also measured bacterial growth of *Atum*GV3101 in the same extracts using gentamicin selection. While no effect on *Atum*GV3101 growth was detected in leaves transiently expressing EV and FLS3 (Figure 3c), transient expression of EFR in *N. benthamiana* reduced *Atum*GV3101 growth (Figure 3c), consistent with a previous study using EFR transgenic plants (Lacombe *et al.*, 2010).

### Depletion of host immunity gene increases bacterial growth in agromonas assay

To test if we could also promote bacterial growth by depleting a host immune component in agromonas assays, we depleted *Nb*FLS2 by RNAi using hairpin (hp) constructs (Yan *et al.*, 2012). We monitored ROS production after flg22 treatment to confirm *Nb*FLS2 depletion. Agroinfiltration of *hpFLS2* had no effect on flg22-induced ROS production three days after agroinfiltration but suppressed the response ten days after agroinfiltration in contrast to *hpGFP* (Figure 4a,b), confirming the selective depletion of *Nb*FLS2.

**Figure 4.**
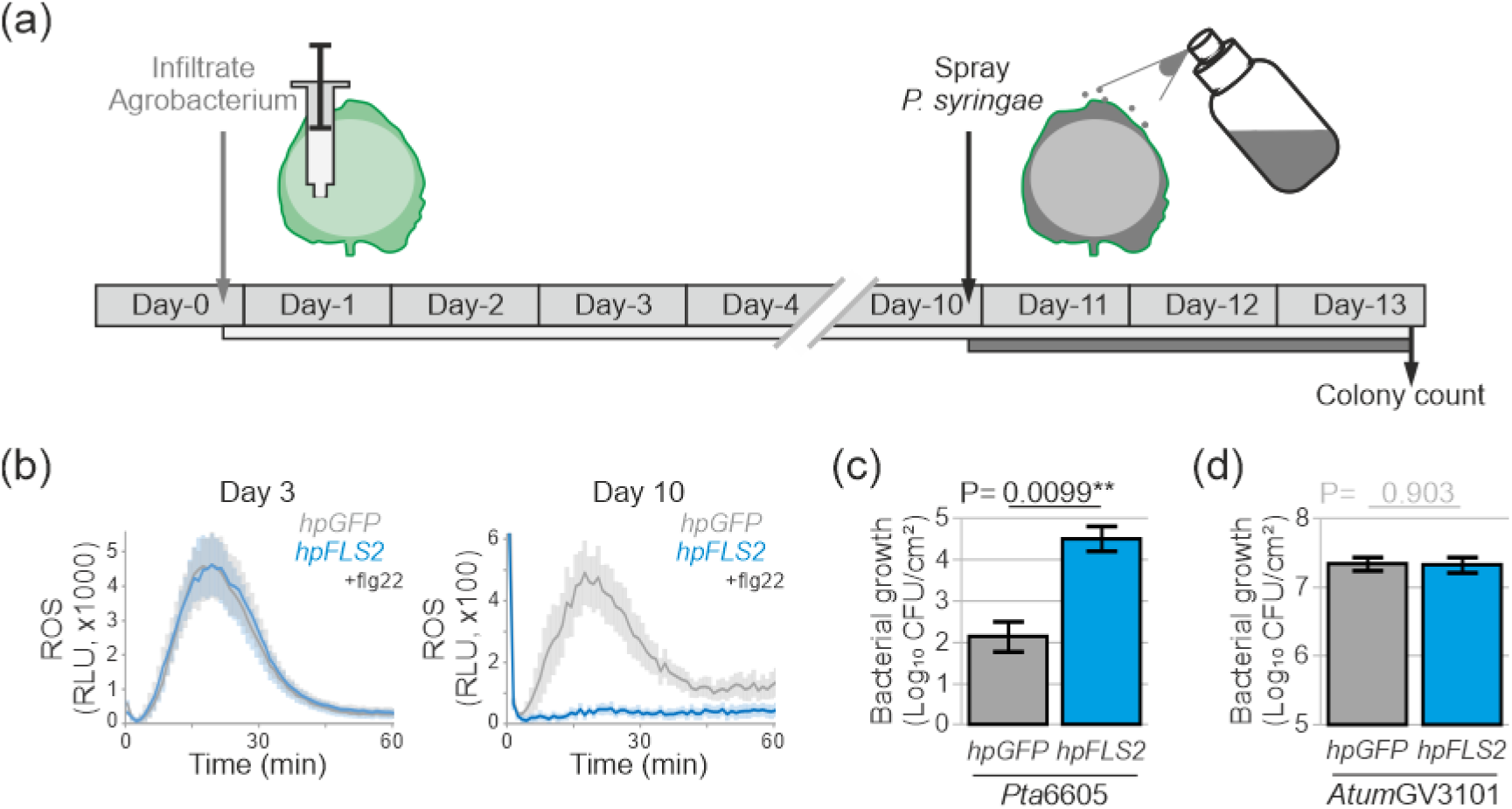
Depletion of host immune gene increases *P. syringae* growth in the agromonas assay. (a) Experimental procedure for studying the role of endogenous immune components in agromonas assay. Ten days after agroinfiltration, agroinfiltrated leaves are infiltrated with *P. syringae* bacteria. Bacterial growth is measured three days later by a classic colony count on LB agar plates containing CFC. (b) FLS2 depletion reduces ROS production upon flg22 treatment. Leaves were agroinfiltrated (OD_600_=0.2) with *hpGFP* (grey) or *hpFLS2* (blue) and at 3 and 10 dpi, leaf discs were treated with 100 nM flg22. Error intervals indicate SE of n=12 replicates. (c) FLS2 depletion increases *P. syringae* growth. Agroinfiltrated leaves expressing *hpGFP* (grey) or *hpFLS2* (blue) were spray-inoculated at 10 dpi with 1×10^8^ CFU/ml *Pta*6605 and bacterial growth was measured three days later using CFC selection. Error bars indicate SE of n=3 biological replicates. Student’s *t*-test statistics (** p<0.01). (d) FLS2 depletion does not affect *Agrobacterium* growth. Bacterial growth of *Atum*GV3101 was measured by plating the leaf extracts described in (c) on medium containing gentamicin. Error bars indicate SE of n=3 replicates. Student’s *t*-test statistics.

We next inoculated these agroinfiltrated leaves depleted for *Nb*FLS2 with *Pta*6605 to measure plant immunity to bacteria. *Nb*FLS2 depletion using *hpFLS2* resulted in significantly more *P. syringae* growth compared to the *hpGFP* control (Figure 4c). These data are consistent with the reported role of *Nb*FLS2 in immunity to *P. syringae* (Segonzac *et al.*, 2011), demonstrating that depletion with hairpin constructs can be used in agromonas assays to study the role of endogenous immune components. By contrast, *Atum*GV3101 grew equally well in both *hpGFP* and *hpFLS2* expressing leaves (Figure 4d), consistent with the absence of immunogenic sequences in flagellin of *A. tumefaciens* that are recognized by *Nb*FLS2 (Hann and Rathjen, 2007).

### Rapid functional analysis of immune components in agromonas assay

To illustrate that the agromonas assay can be used for fast functional analysis (Figure 5a), we generated the non-phosphorylatable mutant EFR^Y836F^ known to be inactive in elf18-induced signalling (Macho *et al.*, 2014). Indeed, agroinfiltrated leaves expressing EFR^Y836F^ were unable to mount an elf18-induced ROS burst, unlike wild-type EFR (Figure 5b), confirming the non-functionality of this EFR^Y836F^ mutant. Consequently, leaves expressing EFR^Y836F^ were more susceptible to *Pto*DC3000(*hQ*), in contrast to leaves expressing wild-type EFR (Figure 5c). Consistent with a role for EFR in conferring resistance to *A. tumefaciens*, growth of *Atum*GV3101 was also reduced by the functional EFR receptor, but not in the presence of EFR^Y836F^ (Figure 5d). These data are consistent with the reported crucial role of Y836 of EFR in immunity to *P. syringae* (Macho *et al.*, 2014), and illustrate that the agromonas assay can be used for quick and robust functional analysis of immune components.

**Figure 5.**
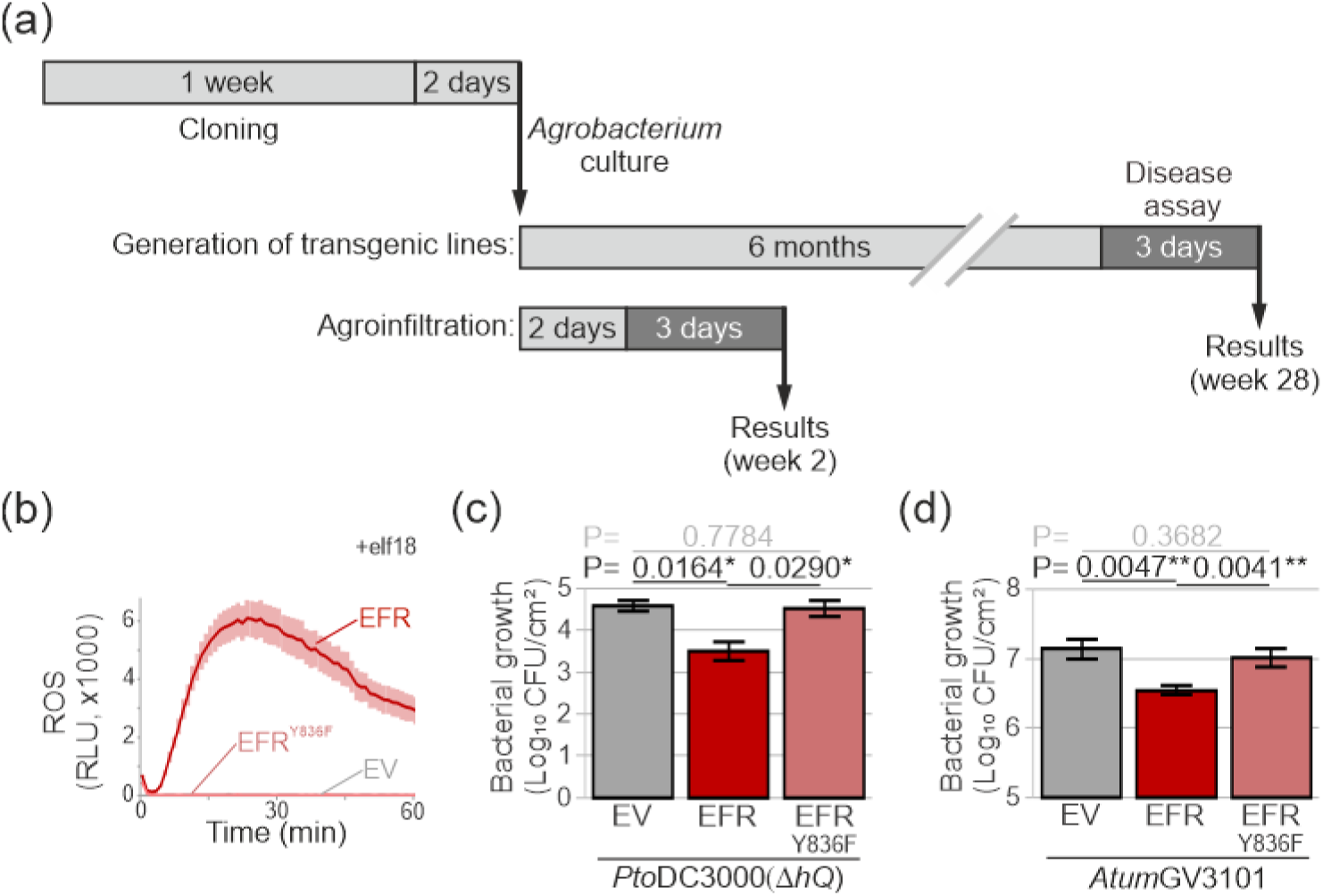
Rapid functional analysis of immune components. (a) Time scale for functional analysis by generation transgenic plants and by agroinfiltration (agromonas assay). (b) Phosphomutant EFR^Y836F^ is unable to trigger ROS burst upon elf18 treatment. Leaves were agroinfiltrated with EFR (red), EFR^Y836F^ (light red) or EV (grey) and the ROS burst was measured at 3 dpi in leaf discs treated with 100 nM elf18. Error intervals indicate SE of n=12 replicates. (c) Phosphomutant EFR^Y836F^ is blocked in elf18-triggered immunity. Two days after agroinfiltration, agroinfiltrated leaves expressing EFR, EFR^Y836F^ or EV were spray-inoculated with 1×10^8^ CFU/ml *Pto*DC3000(∆*hQ*) and bacterial growth was measured three days later using CFC selection. Error bars indicate SE of n=3 biological replicates. Student’s *t*-test statistics (* p<0.05). (d) Agroinfiltration of EFR, but not EFR^Y836F^, reduces Agrobacterium growth. Bacterial growth of *Atum*GV3101 was measured by plating the leaf extracts described in (c) on medium containing gentamicin. Error bars indicate SE of n=3 biological replicates. Student’s *t*-test statistics (** p<0.01).

### T3 effector increases bacterial growth in agromonas assay

To demonstrate that bacterial growth can be increased in agromonas assays by transient expression of microbial effectors, we tested the type III (T3) effector AvrPto, which is a kinase inhibitor blocking PRRs (Xiang *et al.*, 2008). As expected, expression of AvrPto blocked the ROS burst induced by flgII-28 when co-expressed with FLS3 (Figure 6a), consistent with an earlier study (Hind *et al.*, 2016). In addition, AvrPto expression also blocked the ROS burst induced by flg22 (Figure 6b), consistent with an earlier study (Xing *et al.*, 2007). Consequently, agroinfiltration of AvrPto increased growth of *Pto*DC3000(*hQ*) in leaves transiently expressing FLS3 (Figure 6c), demonstrating that agroinfiltration of pathogenic microbial effector supresses host defence. By contrast, *Atum*GV3101 grew equally well on leaves agroinfiltrated with AvrPto (Figure 6d), indicating that AvrPto does not affect *A. tumefaciens* growth by blocking PRRs.

**Figure 6.**
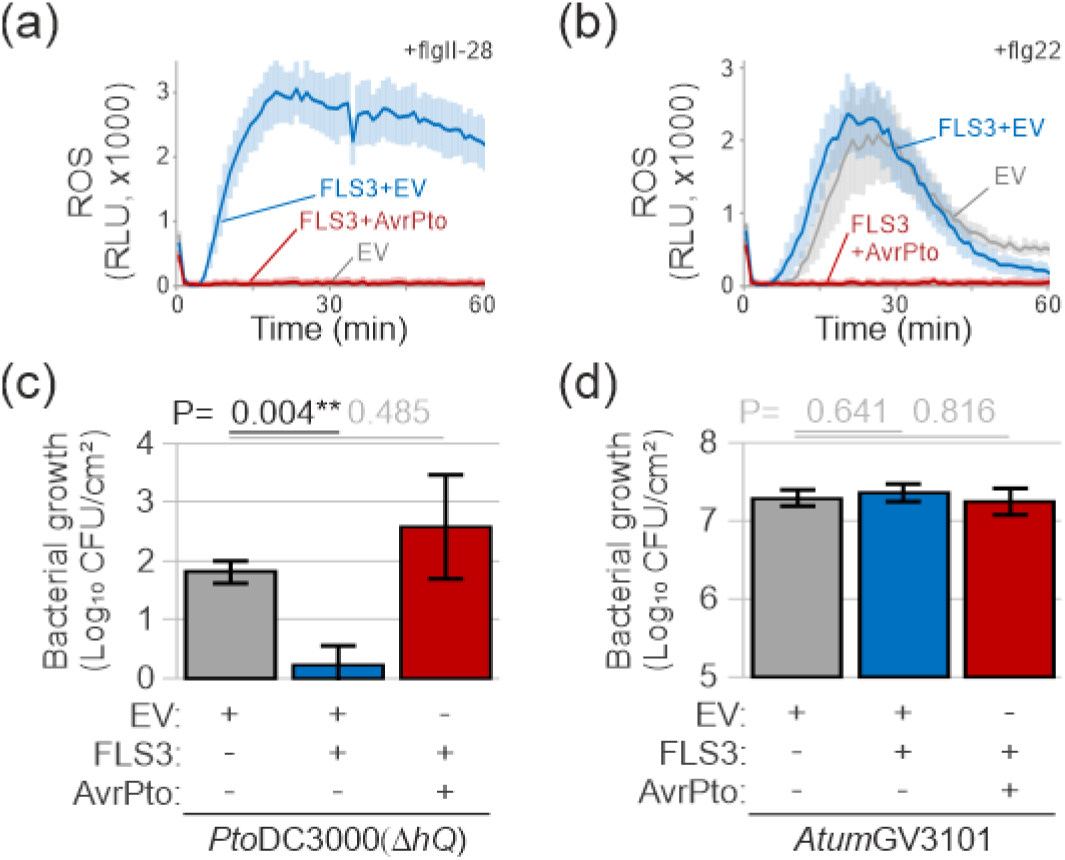
T3 effector supresses immunity in the agromonas assay. (a-b) Expression of AvrPto blocks ROS production upon flgII-28 and flg22 treatment. Leaf discs from agroinfiltrated leaves expressing FLS3 with EV (blue), FLS3 with AvrPto (red) or EV alone (grey) were treated with 100 nM flgII-28 (a) or flg22 (b) and the ROS burst was measured in RLU. Error intervals indicate SE of n=12 replicates. (c) Agroinfiltration of AvrPto increases *P. syringae* growth in *N. benthamiana* and suppresses FLS3-mediated immunity. Two days after agroinfiltration, agroinfiltrated leaves expressing FLS3 in combination with either AvrPto (red) or EV (blue) were spray-inoculated with 1×10^8^ CFU/ml *Pto*DC3000(∆*hQ*) and bacterial growth was measured three days later using CFC selection. Error bars indicate SE of n=3 biological replicates. Student’s *t*-test statistics (** p<0.01). (d) AvrPto does not affect growth of *Atum*GV3101. Bacterial growth of *Atum*GV3101 was measured by plating the leaf extracts described in (c) on medium containing gentamicin. Error bars indicate SE of n=3 biological replicates. Student’s *t*-test statistics.

### Secreted immune hydrolases reduce bacterial growth in agromonas assay

We recently discovered that the β-galactosidase *Nb*BGAL1 contributes to FLS2-mediated immunity by initiating the release of hydrolytic release of flagellin elicitors, presumably by removing the terminal glycan from the flagellin polymer (Buscaill *et al.*, 2019). To test whether immunity triggered by *Nb*BGAL1 can be detected in the agromonas assay, we agroinfiltrated *Nb*BGAL1. Consistent with previous studies (Kriechbaum *et al.*, 2020; Buscaill *et al.*, 2019), apoplastic fluids from leaves of *N. benthamiana bgal1* mutant plants transiently overexpressing *Nb*BGAL1 had strong β-galactosidase activity as *Nb*BGAL1 can cleave galactose from FDG (fluorescein di-β-D-galactopyranoside) and no such activity was detected in the EV control (Figure 7a). These EV and *Nb*BGAL1 expressing leaves were spray-inoculated with *Pta*6605, which carries BGAL1-sensitive glycans. Leaves overexpressing *Nb*BGAL1 had reduced bacterial growth as compared to EV control leaves (Figure 7b), demonstrating that transient expression of *Nb*BGAL1 in *N. benthamiana* increases resistance to *Pta*6605. We also inoculated agroinfiltrated leaves with *Psy*B728a, which carries BGAL1-insensitive glycans. Bacterial growth of *Psy*B728a was not affected by *Nb*BGAL1 when compared to EV expressing leaves (Figure 7c), consistent with the fact that *Nb*BGAL1 acts in immunity only against strains carrying sensitive glycans.

**Figure 7.**
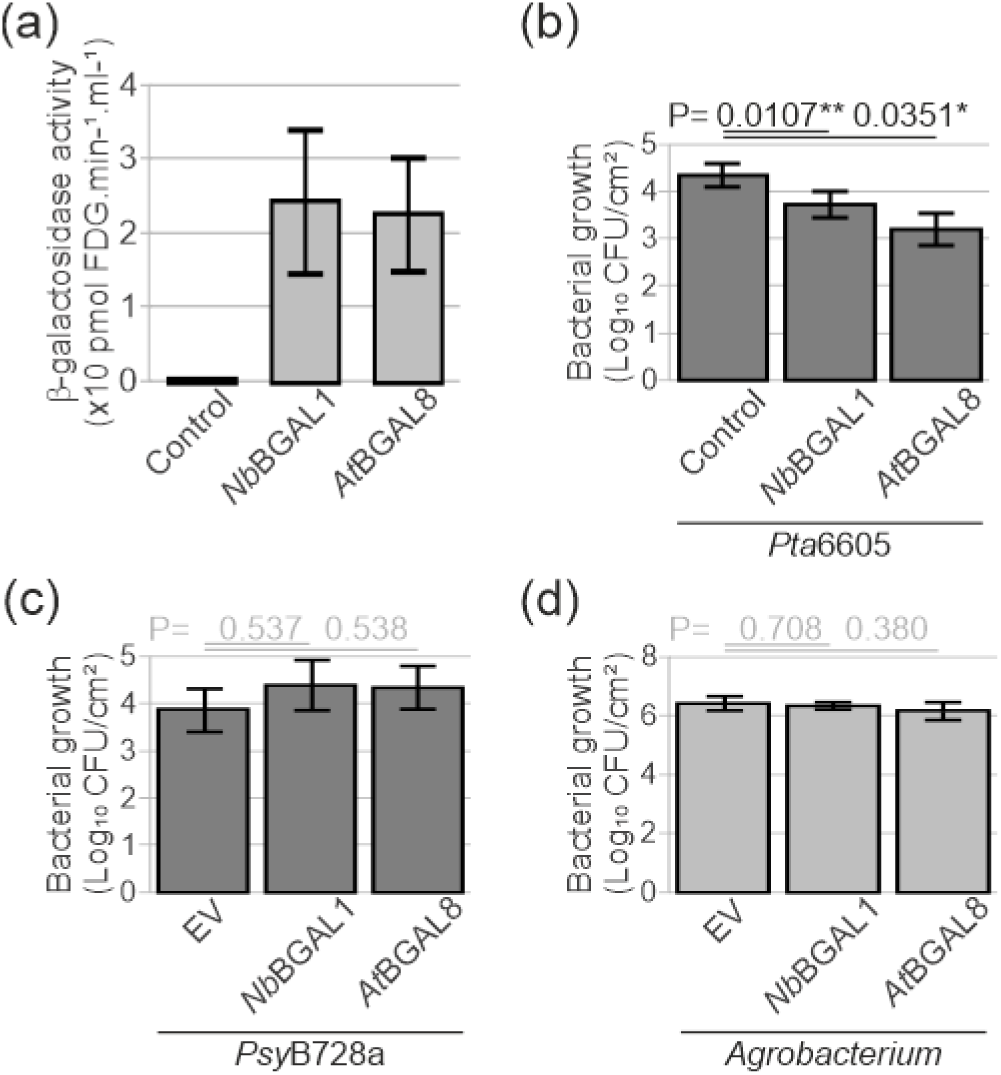
β-galactosidases reduce bacterial growth of BGAL-sensitive strains in agromonas assay. (a) *Nb*BGAL1 and *At*BGAL8 have *β*-galactosidase activity. FDG-hydrolysing activity was measured in apoplastic fluids isolated from *bgal1* mutant leaves transiently expressing *Nb*BGAL1 or *At*BGAL8. Error bars indicate SE of n=3 biological replicates. (b) Agroinfiltration of *Nb*BGAL1 and *At*BGAL8 reduce *Pta*6605 growth. Two days after agroinfiltration, agroinfiltrated leaves expressing *Nb*BGAL1 or *At*BGAL8 were spray-inoculated with 1×10^8^ CFU/ml *Pta*6605 and bacterial growth was measured three days later using CFC selection. Error bars indicate SE of n=6 biological replicates; *t-*test P values (* p<0.05). (c) *Nb*BGAL1 or *At*BGAL8 do not reduce *Psy*B728a growth. Two days after agroinfiltration, agroinfiltrated leaves expressing *Nb*BGAL1 or *At*BGAL8 were spray-inoculated with 1×10^8^ CFU/ml *Psy*B728a and bacterial growth was measured three days later using CFC selection. Error bars indicate SE of n=3 biological replicates; *t-*test P values. (d) *Nb*BGAL1 and *At*BGAL8 do not affect *Agrobacterium* growth. Bacterial growth of *Atum*GV3101 was measured by plating the leaf extracts described in (c) on medium containing gentamicin. Error bars indicate SE of n=3 biological replicates. Student’s *t*-test statistics.

*Atum*GV3101 grew equally well on both plants agroinfiltrated with *Nb*BGAL1 and EV (Figure 7d), indicating that *Nb*BGAL1 does not affect *Agrobacterium* growth, consistent with the absence of immunogenic flagellin peptides triggering *Nb*FLS2 (Hann and Rathjen, 2007).

We finally tested whether the *Nb*BGAL1 ortholog in Arabidopsis, *At*BGAL8 can provide immunity to strains carrying *Nb*BGAL1-sensitive glycans. *Nb*BGAL1 and *At*BGAL8 share 72% amino acid identity and *At*BGAL8 carries the catalytic residues (Figure S2). Similarly to *Nb*BGAL1, *At*BGAL8 has *β*-galactosidase activity when produced by agroinfiltration in *N. benthamiana bgal1* mutant plants (Figure 7a) and as with *Nb*BGAL1, agroinfiltration of *At*BGAL8 reduces growth of *Pta*6605 (Figure 7b) but did not affect the growth of *Psy*B728a (Figure 7c). Thus, similarly to *Nb*BGAL1, *At*BGAL8 can confer specific immunity to *P. syringae* strains carrying BGAL1-sensitive glycans. By contrast, *Atum*GV3101 grew equally well on agroinfiltrated leaves expressing *At*BGAL8 (Figure 7d), indicating that *At*BGAL8 does not affect *Agrobacterium* growth. These data demonstrate that the agromonas assay can be used to study diverse components of the immune system.

## DISCUSSION

Agromonas is an easy, robust, and simple assay to monitor *P. syringae* growth in agroinfiltrated leaf tissues both upon infiltration and spray-inoculation of *P. syringae*. Using well established PAMP receptors, a T3 effector and immune hydrolases, we demonstrated that the agromonas assay can be used to study components that enhance immunity (*e.g.* FLS3, EFR and BGAL1) and reduce immunity (*e.g. hpFLS2* and AvrPto). This assay is now routinely used in our lab to study various novel components of plant immunity, perform structure-function analysis, and study putative roles of effectors in immune suppression. This manuscript sets examples and parameters for this assay so it can be used widely by the research community.

### Four practical considerations to take away

This manuscript establishes the methodology of the agromonas assays. As for all disease assays, experimental conditions are of fundamental importance. There are four essential parameters that should be considered for agromonas assays.

First, it is essential to use EV and *hpGFP* controls in overexpression and depletion assays, respectively. Agrobacterium suppresses *P. syringae* growth directly and/or indirectly, so a mock control (*i.e.* buffer infiltration) is not very useful. The control should be based on the same Agrobacterium strain carrying the same vector and at same final OD_600_.

Second, it is worth simultaneously monitoring growth of Agrobacterium simply using gentamicin selection, when the *P. syringae* strain being used is not gentamicin resistant. Occasionally, Agrobacterium growth is also affected by modulation of host immunity and this may affect protein expression. However, in the cases described here, Agrobacterium growth was reduced by EFR consistent with the literature (Kunze *et al.*, 2004), but immunity to *P. syringae* was still enhanced, indicating that *P. syringae* growth was suppressed by EFR not by reduced Agrobacterium levels.

Third, it is crucial to allow sufficient protein accumulation in the agroinfiltrated leaves prior *P. syringae* infection. This may differ between proteins. Likewise, depletion of endogenous proteins by *hp*RNAi may need time. For instance, three days upon agroinfiltration of *hpFLS2* was not enough to deplete endogenous *Nb*FLS2.

Fourth, we recommend analysing at least 3 (ideally 6) plants per condition per experiment and repeating experiments at least 3 times. This is normal practice in *P. syringae* infections and will undoubtedly display the high reproducibility of the agromonas assay.

### Five limitations of agromonas assays

Despite the broad versatility of the agromonas assay, we would like to point out several limitations of the assay that should be considered for future experiments.

First, excessively high expression levels could cause artefacts, but this problem can be mitigated by using different promoters (Grefen *et al.*, 2010). We like to note that many microscopy studies on fluorescent tagged proteins show the expected subcellular localisations (Bally *et al.*, 2018), indicating that *N. benthamiana* usually delivers proteins at the intended site.

Second, *A. tumefaciens* strains employed for agroinfiltration are non-oncogenic but they still trigger host immunity. Indeed, the receptor CORE (cold shock protein receptor) of *N. benthamiana*, tomato (*S. lycopersicum*), tobacco (*N. tabacum*) and potato (*S. tuberosum*) recognizes the csp22 epitope of CSP from *A. tumefaciens* (Felix and Boller, 2003; Wang *et al.*, 2016). Agroinfiltration into *N. tabacum* leaves elicits a low level of callose deposition (Rico *et al.*, 2010). Increased host immunity is visible in reduced *P. syringae* growth and may be a limitation to study specific immune components.

Third, some immune responses are supressed by agroinfiltration. For instance, agroinfiltration into *N. tabacum* leaves results in reduced abscisic acid (ABA) levels and salicylic acid (SA) production (Rico *et al*., 2010).

Fourth, despite the fact that *N. benthamiana* is commonly used as a model for plant-pathogen interactions (Goodin *et al.*, 2008) and possesses PRR co-receptors (Heese *et al.*, 2007; Liebrand *et al.*, 2013), it may not possess all supporting components when testing immune components from other plant species. For example, *N. benthamiana* possesses ZAR1, but lacks ZED1 for the recognition of the bacterial effector HopZ1 (Baudin *et al.*, 2017).

Fifth, quantification of bacterial growth by colony count assays remains a bottle neck for high-throughput screening. Alternative pathogen infection assays have been developed to increase throughput. For instance, bacterial DNA can be quantified by real-time PCR (Ross and Somssich, 2016), but this technique does not distinguish between living and dead bacteria, overestimating the titres of living bacteria (Rooney *et al.*, 2020). An alternative approach for monitoring bacterial density is using bioluminescence (Fan, Crooks and Lamb, 2008), but this method requires the transformation of each bacterial strain with the *luxCDABE* operon (Fan, Crooks and Lamb, 2008). Also, bioluminescence reflects the metabolite state rather than bacterial viability and cannot be used to detect low titres. However, bioluminescence might be a powerful approach to increase throughput in specific contexts.

### Six opportunities of using agromonas assays

We believe that the agromonas assay can be applied to a wide range of applications. There are at least five main opportunities for a wider application of the agromonas assay.

First, the agromonas assay is a rapid and easy method to subject plant and pathogen proteins in immunity for structure-function analysis, without the need of transgenic plants. For instance, we tested the non-phosphorylatable mutant EFR^Y836F^ (Macho *et al.*, 2014) to demonstrate that agromonas assay facilitates rapid structure-function analysis of immune components.

Second, since CFC selects *Pseudomonas* spp., the agromonas assay works without genetic manipulation of *P. syringae*, allowing the study of the available repertoire of *P. syringae* mutants and strains. For instance, *Pto*DC3000 polymutants (Wei *et al.*, 2015) and *Psy*B728a mutants (Vinatzer *et al.*, 2006) can be tested in agromonas assays.

Third, although in this study we only used *N. benthamiana*, we anticipate that the agromonas assay can be easily adapted to plant species suitable for agroinfiltration such as potato (Du *et al.*, 2014), tobacco (Van der Hoorn *et al.*, 2000), pea (Guy *et al.*, 2016), *Medicago* (Picard *et al.*, 2013), grapefruit (Figueiredo *et al.*, 2011), lettuce (Chen *et al.*, 2016), tomato (Wroblewski *et al.*, 2009), flax (Dodds *et al.*, 2006), cassava (Zeng *et al.*, 2019), strawberry (Guidarelli and Baraldi, 2015) and *Mucuna bracteate* (Abd-Aziz *et al.*, 2020). This facilitates studies of immunity in other plant species using corresponding *P. syringae* pathovars.

Fourth, for plant species that are not amendable to agroinfiltration, *N. benthamiana* remains an excellent heterologous expression system to study proteins from various organisms (plants, microbes and animals). We demonstrate this by studying EFR and *At*BGAL8 from Arabidopsis and FLS3 from tomato. We anticipate that agromonas assays can be used to study more immune-related genes or QTL (quantitative trait locus) from various plants and microbes. For instance, PAMPs receptors LORE and LYM1/3 (Ranf *et al.*, 2015; Willmann *et al.*, 2011), are absent in *N. benthamiana* and should reduce bacterial growth in agromonas assays.

Fifth, *P. syringae* growth can be monitored in agromonas assay using both infiltration and spray-inoculation, testing both post- and pre-invasive immunity. Here, we used the method of inoculation described in literature for each immune component tested (Macho *et al.*, 2014; Segonzac *et al.*, 2011; Xiang *et al.*, 2008; Buscaill *et al.*, 2019). We anticipate that leaf-dipping and vacuum inoculation should also be applicable in agromonas assays.

Sixth, agromonas assays can be used to study depletion of host immunity genes. We demonstrated this by depleting *Nb*FLS2 using *hpFLS2*, which resulted in a significant enhancement of *P. syringae* growth. Similar results were observed with depletion of *Nb*FLS2 by VIGS (virus-induced gene silencing) (Segonzac *et al.*, 2011). However, VIGS is at least four weeks slower than *hp*RNAi technique and VIGS requires approval for work with modified TRV (tobacco rattle virus). We anticipate that agromonas assays can also be used to study the depletion of additional endogenous immune components, such as MAPKs (mitogen-activated protein kinases), CDPKs (calcium-dependent protein kinases), transcription factors, PAMP receptors and NLRs (nucleotide-binding/leucine-rich repeat). However, depletion of these components with transient *hp*RNAi will depend on the stability of the protein.

In conclusion, the ability to characterise immune components from plants and microbes using agromonas assays will speed up our understanding of plant immune system and microbial colonisation and generate promising new strategies for crop protection.

## EXPERIMENTAL PROCEDURES

### Plants

*Nicotiana benthamiana* plants were grown in a growth chamber at 21 °C and ~60% relative humidity with a 16 h photoperiod and a light intensity of 2000 cd.sr.m^−2^.

### Molecular cloning

All constructs were generated using standard molecular biology procedures. All vectors used in this study are listed in Table 1. EFR (At5g20480) and *At*BGAL8 (At2g28470) were amplified from *A. thaliana* ecotype Col-0 complementary DNA (cDNA), and *Sl*FLS3 (LOC101248095/Solyc04g009640) was amplified from *S. lycopersicum* cv. Rio Grande genomic DNA (gDNA) using primers listed in Table 2. *BGAL1* was amplified from *N. benthaminana* cDNA using primers listed in Table 2. The PCR products were combined with pICH51288 (Engler *et al.*, 2014), pICH41414 (Engler *et al.*, 2014), and pJK001c (Paulus *et al.*, 2020) in a BsaI GoldenGate reaction to generate pPB069 (pL2M-2×35S::EFR), pJK668 (pL2M-2×35S::FLS3), pJK646 (pL2M-2×35S::*Nb*BGAL1) and pJK645 (pL2M-2×35S::*At*BGAL8), respectively. The EFR tyrosine mutant EFR^Y836F^ was generated by site directed mutagenesis using primers listed in Table 3. All binary plasmids were transformed into *Agrobacterium tumefaciens* GV3101 (pMP90) by freeze-thawing and transformants were selected by kanamycin resistance.

**Table 1.** Binary vectors generated in this study.

| Name | Description |
| --- | --- |
| pPB069 | 2x35S::EFR |
| pJK668 | 2x35S::FLS3 |
| pPB076 | 2x35S::EFR<sup>Y836F</sup> |
| pJK646 | 2x35S::NbBGAL1 |
| pJK665 | 2x35S::AtBGAL8 |
| pPB070 | hp::GFP |
| pPB072 | hp::NbFLS2 |

**Table 2.**
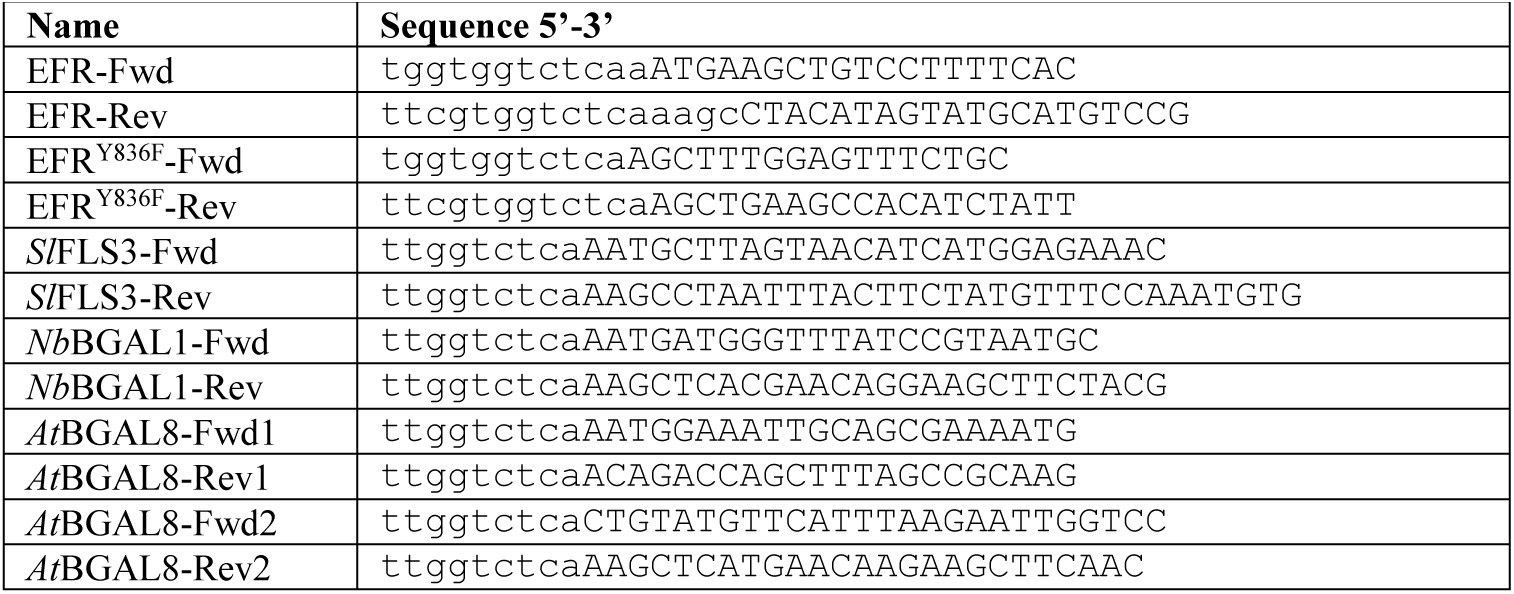
Primers used in this study.

**Table 3.** Bacterial strains used in this study

| Strain | Reference |
| --- | --- |
| *Pseudomonas syringae* pv. *tomato* DC3000($\Delta hQ$) | Wei *et al.*, 2007 |
| *Pseudomonas syringae* pv. *tabaci* 6605 | Taguchi *et al.*, 2006 |
| *Pseudomonas syringae* pv. *syringae* B728a | Vinatzner *et al.*, 2006 |
| *Pseudomonas syringae* pv. *aptata* NCPPB 3539 | Higgins, NCPPB |
| *Agrobacterium tumefaciens* GV3101 | Koncz and Schell 1986 |
| *Xanthomonas campestris* pv. *campestris* 8004 | NCBI:txid314565 / Qian *et al.*, 2005 |
| *Xanthomonas campestris* pv. *vesicatoria* 85-10 | NCBI:txid316273 / Thieme *et al.*, 2005 |
| *Xanthomonas axonopodis* pv. *citri* 306 | NCBI:txid190486 / da Silva *et al.*, 2002 |
| *Xanthomonas a.c.* pv. *citri* | NCPBP 3233 |
| *Xanthomonas c.c.* pv. *campestris* | NCPBP528 |
| *Xanthomonas t.* pv. *translucens* | NCPBP 973 |

### Silencing by RNA interference (RNAi)

An intron-containing hairpin RNA (ihpRNA) construct targeting a conserved region in *FLS2a/b* was designed to silence both *FLS2* homologs (*i.e.* NbD013936.1 and NbD024362.1) detected in the NbDE database (Kourelis *et al.*, 2019). The 300 bp fragment of *GFP* (Sequence A) and the 300 bp fragment of *FLS2a/b* (Sequence B) used for RNAi were commercially synthesized (Invitrogen, Carlsbad, CA, USA). The fragments were cloned into the pRNAiGG vector (Yan *et al.*, 2012) using *BsaI* restriction sites resulting in vector pPB070 and pPB072, respectively (Table 2). The binary constructs were transformed into *Atum*GV3101 (pMP90) by freeze-thawing and transformants were selected by kanamycin resistance. Three-week-old *N. benthamiana* leaves were agroinfiltrated with the hairpin silencing construct at a final OD_600_ = 0.2. Further experiments (ROS production and spray infection) were performed ten days after agroinfiltration.

#### Sequence A

fragments used for silencing *GFP*. <u>Underlined</u>: BsaI restriction sites.

acca<u>ggtct</u>caggagATGCGTAAAGGCGAAGAGCTGTT CACTGGTGTCGTCCCTATTCTGGTGGAACTGGATGGTGATGTCAACGGTCATAAGTTTTCCGTGCGTGGCGAGGGTGAAGGTGACGCAACTAATGGTAAACTGACGCTGAAGTT CATCTGTACTACTGGTAAACTGCCGGTACCTTGGCCGACTCTGGTAACGACGCTGACTTATGGTGTTCAGTGCTTT GCTCGTTATCCGGACCATATGAAGCAGCATGACTTCTTCAAGTCCGCCATGCCGGAAGGCTATGTGCAGGAACGCA CGATTTCCTTTacgat<u>gagacct</u>ggt

#### Sequence B

fragments used for silencing *FLS2*. <u>Underlined</u>: BsaI restriction sites.

acca<u>ggtctc</u>aggagGGAACGATCCCAGATGAGATTGG CGTGTTAGAAATGGTTCAAGAGATTGATATGTCAAACA ACAATCTATCAGGCAGCATTCCCAGTTCCTTTGGACGA TGCAAAAACTTATTCTCACTGGACCTATCAGGAAACAT GCTCTCTGGTCCTGCTCCAGGTGCAGTTCTTACCAAAT TAAGTGAGCTTGTGTTTTTGAATCTCTCAAGGAACAGA TTAGAAAGCGAACTTCCTGAAATGGCAGGATTGCCACA TCTTCGCTCTGTTGATCTTTCACATAACAAGTTCAAGG GAATCATTCCAacgat<u>gagacct</u>ggt

### Agroinfiltation

For transient expression of proteins in *N. benthamiana*, overnight cultures of *A. tumefaciens* GV3101 (*Atum*GV3101) carrying binary vectors were harvested by centrifugation. Cells were resuspended in induction buffer (10 mM MgCl_2_, 10 mM MES pH5.0, and 100 µM acetosyringone) and mixed (1:1) with bacteria carrying silencing inhibitor P19 at OD_600_ = 0.5. After 1 h at 21 °C, cells were infiltrated into three leaves of 4-week-old *N. benthamiana*. Leaves were harvested and processed at the indicated days after agroinfiltration.

### Bacterial strains

The bacterial strains used in this study are listed in Table 3. *Pseudomonas* and *Xanthomonas* strains were grown in Luria-Bertani (LB) medium at 28 °C. For the infection assays, bacteria were cultured in LB medium containing 10 mM MgCl_2_ at 28 °C. *Agrobacterium tumefaciens* strains were grown in LB medium containing 50 μg/ml rifampicin, 10 μg/ml gentamicin and 50 μg/ml kanamycin at 28 °C.

### Oxidative burst assays

The generation of reactive oxygen species (ROS) was measured by a luminol-based assay on leaf discs adapted from (Smith *et al*., 2014). Luminol (Sigma-Aldrich, Saint Louis, MO, USA) was dissolved in dimethyl sulfoxide at a concentration of 10 mg/ml and kept in the dark. Horseradish peroxidase (Thermo Fisher Scientific, Waltham, MA, USA) was dissolved in water at a concentration of 10 mg/ml. Leaf discs (6 mm diameter), were incubated overnight in water in petri dishes. Leaf discs from agroinfiltrated leaves were deposed in a 96-well plate, one leaf disc per well (Costar, Kennebunk, ME, USA). 100 µl of 25 ng/µl luminol, 25 ng/µl HRP and 100 nM elf18 (Kunze *et al*., 2004) or 100 nM flg22 (Felix *et al*., 1999) or 100 nM flgII-28 (Cai *et al*., 2011) was added and chemiluminescence was recorded immediately in relative light unit (RLU) using an Infinite M200 plate reader (Tecan, Mannedorf, Switzerland). Measurements were taken every minute for 1 h. Standard errors were calculated at each time point and for each treatment. Experiments were repeated at least three times.

### Bacterial growth upon inoculation

For syringe-inoculations, an overnight culture was washed and resuspended in sterile water to a density of 1×10^6^ CFU/ml and infiltrated into agroinfiltrated leaves using a blunt syringe via the abaxial side of the leaves. For spray-inoculations, an overnight culture was washed and resuspended in sterile water to a density of 1×10^8^ CFU/ml and sprayed onto adaxial surfaces of agroinfiltrated leaves. Before inoculation, plants were covered with a humidified dome for one day. After infection, plants were re-covered with the dome and kept for three days in a growth cabinet at 21 °C. For determination of *in planta* bacterial growth, three leaf discs (1 cm diameter) were excised three days after inoculation from inoculated leaves. Each leaf disc was soaked in 15% H_2_O_2_ for 2 min to sterilise the leaf surface. Leaf discs were washed twice in sterile water and dried under sterile condition for 30 min. Leaf discs were then ground in sterile water for 5 min using the tissue-lyser and metal beads (Biospec Products, Bartelsville, OK, USA). Serial dilutions of the homogenate were plated onto LB agar supplemented with either gentamicin (10 µg/ml) for selection of *Atum*GV3101 or CFC (Oxoid™ C-F-C Supplement) at 1x concentration, prepared according to the manufacturer’s instructions for selection of *P. syringae* strains and incubated at 28 °C. Colonies were counted after 36 h incubation at 28 °C. The p-value was calculated using the two-tailed Student’s *t*-test to binary compare bacterial growth between agroinfiltrated plants.

### Statistics

All values shown are mean values and the error intervals shown represent standard error of the mean (SE), unless otherwise indicated. P-values were calculated using the two-tailed Student’s *t*-test. All experiments have been reproduced and representative datasets are shown.

### Protein alignment

Sequences were aligned using Clustal Omega (Sievers *et al.*, 2011). Alignment was visualised and analysed using Jalview (Waterhouse *et al.*, 2009) and edited using CorelDraw (Corel Corporation, Ottawa, Ontario, Canada).

## ACKNOWLEDGEMENTS

We thank Alan Collmer for providing the *Pto*DC3000(∆*hQ*) mutant; Yuki Ichinose for providing the *Pta*6605 strain; Tolga Bozkurt for providing the pRNAiGG vector; Nicola J. Patron and Sylvestre Marillonnet for providing Golden Gate plasmids; Ursula Pyzio for plant care; Sarah Rodgers and Caroline O’Brien for technical support.

## Author contributions

PB, NS, GP and RH conceived the project; PB, NS, YL and JK performed experiments; PB and RH wrote the manuscript with input from all authors.

## Data and materials availability

all data is available in the manuscript, the supplementary materials, and the cited references.

## SUPPORTING INFORMATION

**Table S1.** Selection of bacterial strains on medium supplemented with CFC or gentamicin.

| Strain | CFC | Gentamicin |
| --- | --- | --- |
| *Pseudomonas syringae* pv. *tomato* DC3000($\Delta hQ$) | Growth | No growth |
| *Pseudomonas syringae* pv. *tabaci* 6605 | Growth | No growth |
| *Pseudomonas syringae* pv. *syringae* B728a | Growth | No growth |
| *Pseudomonas syringae* pv. *aptata* NCPPB 3539 | Growth | No growth |
| *Xanthomonas campestris* pv. *campestris* 8004 | No growth | No growth |
| *Xanthomonas campestris* pv. *vesicatoria* 85-10 | No growth | No growth |
| *Xanthomonas axonopodis* pv. *citri* 306 | No growth | No growth |
| *Xanthomonas a.c.* pv. *citri* NCPPB 3233 | No growth | No growth |
| *Xanthomonas c.c.* pv. *campestris* NCPPB528 | No growth | No growth |
| *Xanthomonas t.* pv. *translucens* NCPPB 973 | No growth | No growth |
| *Agrobacterium tumefaciens* GV3101 | No growth | Growth |
| *Agrobacterium tumefaciens* C58C1 | No growth | Growth |

**Figure S1.**
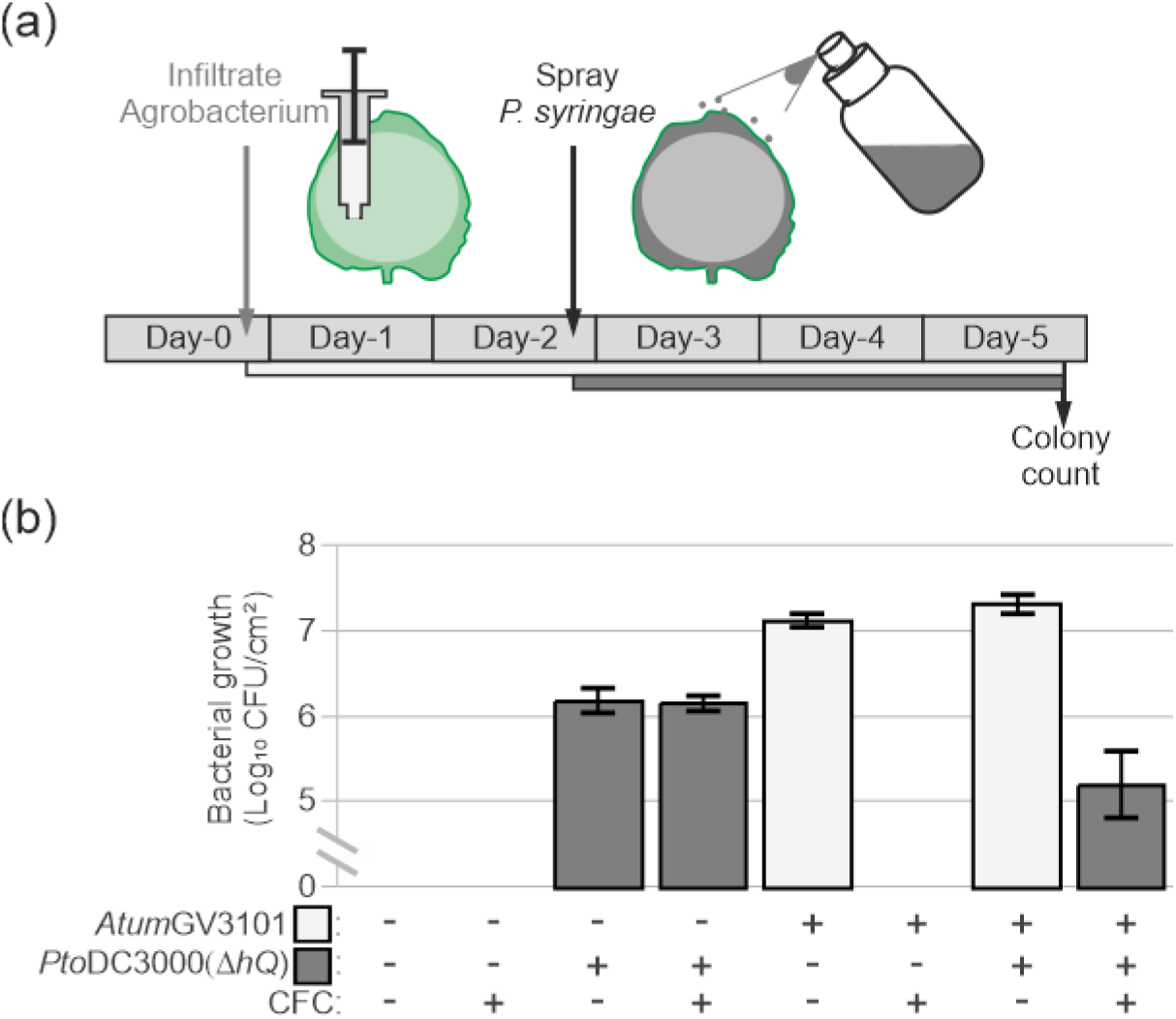
Concept of Agromonas method. (a) Experimental procedure for Agromonas assay to monitor *P. syringae* growth in agroinfiltrated leaves. Two days after agroinfiltration, leaves are spray-infected with *P. syringae*. Three days later, bacterial growth is measured with a classical colony count on medium containing CFC. (b) Selective isolation of *Pto*DC3000(*hQ*) from agroinfiltrated leaves. Agroinfiltrated leaves were spray-infected with 1×10^8^ CFU/ml *Pto*DC3000(*hQ*) and three days later, *Pto*DC3000(*hQ*) and *Atum*GV3101 bacteria were selected on LB agar plate with and without CFC, respectively. CFU, colony forming unit.

**Figure S2.**
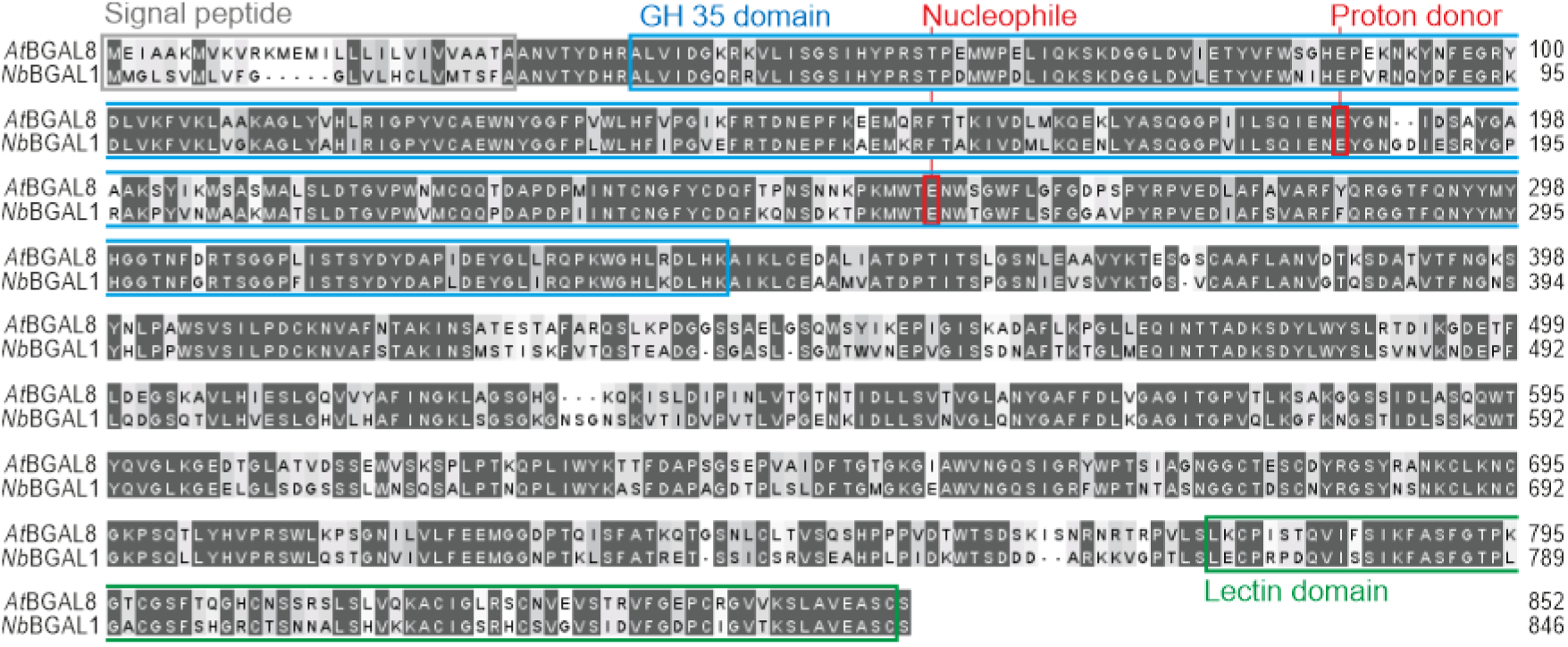
Alignment of *Nb*BGAL1 and *At*BGAL8 protein sequences. Clustal Omega alignment of *Nb*BGAL1 and *At*BGAL8. Amino acid residues are shaded black if identical or grey if similar. Signal peptide sequences (grey) were extracted from the UniProtKB database and also predicted by SignalP 5.0 server. The glycoside hydrolase 35 domain (GH35, blue) contains two active catalytic glutamate residues (nucleophile and proton donor, red). Predicted lectin domain (green). Annotations were extracted from the Pfam database.

